# Spatial Single-Cell Proteomics Reveals Molecular Trajectories Of Tangle-Bearing Neurons In Alzheimer’s Disease

**DOI:** 10.64898/2026.04.26.720932

**Authors:** M.S. Foiani, M. Bourdenx, L. Kraller, R.S. Nirujogi, A. Yiu, H. Davies, S. Patel, L.S. Damoc, L. Mitchener, Z. Jaunmuktane, F. Coscia, K.E. Duff

**Author notes:** These authors contributed equally (M.S. Foiani, M. Bourdenx, L. Kraller).

## Abstract

Neurofibrillary tangles composed of hyperphosphorylated tau are a defining pathological hallmark of Alzheimer’s disease (AD); however, the pathways and mechanisms associated with the transition from physiological tau to tangle pathology remain unclear. Here, we integrate laser microdissection of post-mortem, fixed human AD brain tissue labelled with an antibody recognizing tangle-associated phospho-tau (AT8) with mass spectrometry–based proteomics, applied to individual neurons and to small neuronal pools. This approach identified ∼2,000 and ∼5,000 proteins, respectively, and enabled direct detection of disease-associated tau phosphorylation sites without prior enrichment. A layered analysis of the proteome of tangle-positive and tangle-negative neurons revealed heterogeneous disease-associated states. Pseudotime analysis, combined with an AI-driven analytical framework, indicates that neurons do not segregate into discrete classes but instead organize along a continuum of proteomic changes that correlate with tau abundance. This organization enabled the construction of a trajectory of pathological neuronal responses that can be resolved within an individual brain. Early stages of this trajectory are characterized by coordinated remodeling of proteostasis networks, including reduced proteasome component abundance and increased lysosomal acidification machinery, followed by disruption of synaptic pathways. Notably, despite extensive proteomic remodeling, neurons bearing tangles show little evidence of activated cell-death programs, suggesting prolonged molecular adaptation rather than acute degeneration. Together, these findings establish a framework for single-cell–resolved proteome analysis of human brain disease *in situ* and define a continuum of neuronal states underlying tau pathogenesis, revealing early vulnerabilities and adaptive responses during AD progression.

## Main

Alzheimer’s disease (AD), the most common cause of dementia, is characterized by the extracellular accumulation of amyloid-β plaques and intraneuronal aggregates of hyperphosphorylated tau into neurofibrillary tangles (NFTs). While both pathologies define the disease, the abundance of NFTs more closely correlates with cognitive decline (Gómez-Isla et al. 1997). However, recent studies in human AD brains suggest a disconnect between NFT accumulation and death of the affected neuron (Zwang et al. 2024). These findings highlight an important gap in our understanding of the molecular consequences of tangle formation in the human brain and how neuronal integrity is affected during this process.

Despite the broad and constitutive expression of tau throughout the brain, only a subset of neurons accumulate NFTs, while neighboring neurons remain unaffected (Braak and Braak 1991). This phenotypic variability allows for a direct, within-brain comparison of the response to neurons across a continuum of disease states.

To investigate the molecular consequences of tau pathology at the single-cell level relative to neighboring spared neurons, we applied exploratory mass spectrometry (MS)-based spatial proteomics to individual neurons laser microdissected from formalin-fixed, paraffin-embedded (FFPE) prefrontal cortex (BA9) layer II tissue from AD cases at Braak stage VI (**Table 1**). Neurons were stratified according to pTau-status: “pTau positive” neurons were identified by immunolabeling with the AT8 antibody recognizing phosphorylated epitopes that accumulate in NFTs, whereas “pTau negative” neurons were defined by counterstaining (cresyl-violet) and absence of AT8 signal. We collected both small pools of 20 neurons (hereafter “mini-pool”; n=2-4 pools per condition and cases, total n=44 over 10 cases) and individual neurons (7µm sections of single-cell bodies; n=19-46 cells per patient, total n=187 over 5 cases). Mini-pools allow for higher proteome coverage and quantitative robustness, whereas single-cell proteomes provide higher resolution of neuron-to-neuron differences. All samples were collected and processed using loss-reduced tissue proteomics workflow for ultra-low input of FFPE material (**Fig. 1a**) (Makhmut et al. 2023). Proteomes were acquired on a timsTOF Ultra 2 mass spectrometer and analyzed with DIA-NN (Demichev et al. 2020), with a predicted human spectral library.

**Figure 1.**
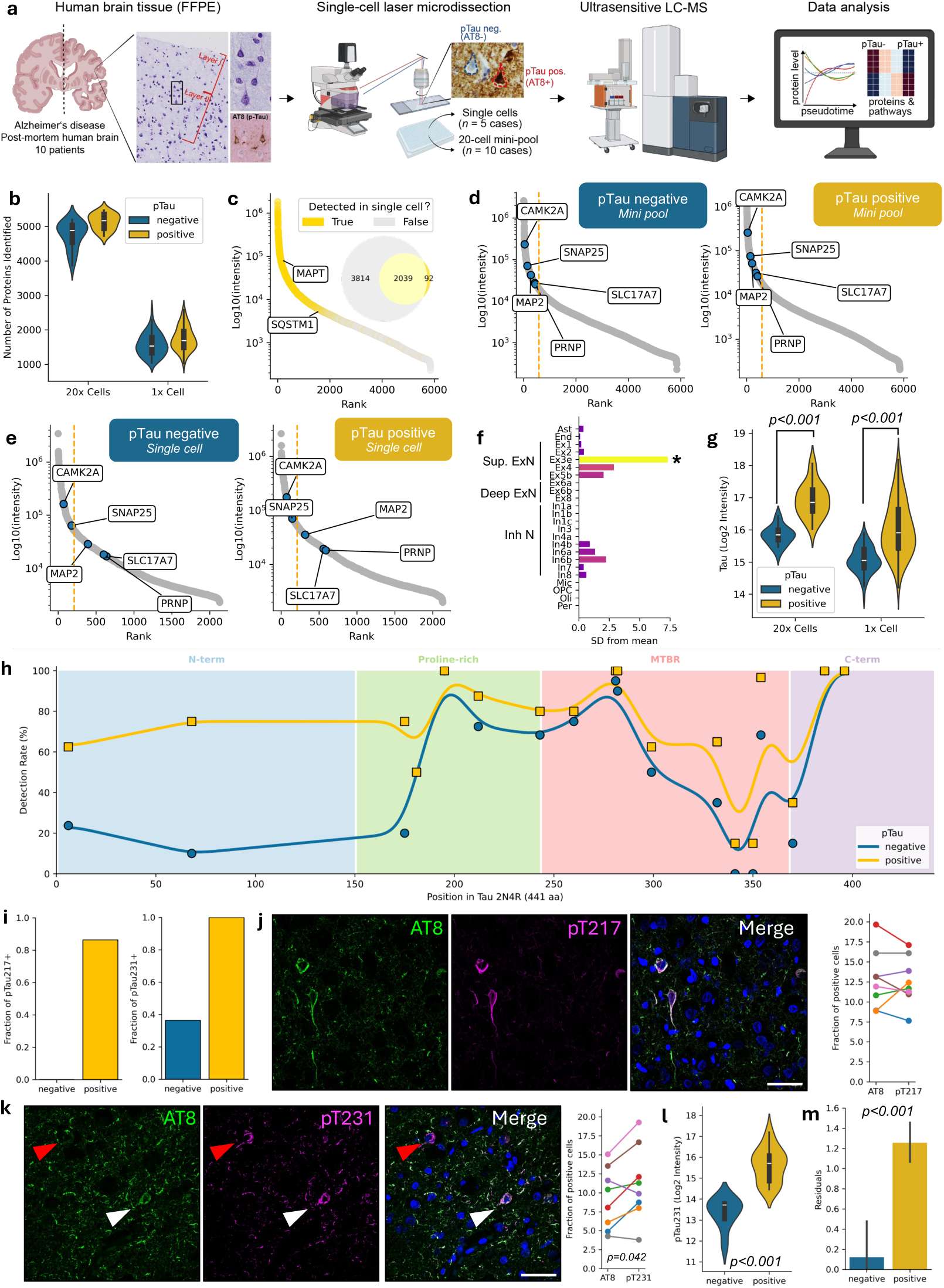
Single-cell proteomics of tangle-bearing neurons. (**a**) Graphical summary of data collection and acquisition. (**b**) Number of detected proteins from 20x cells (mini-pool) and single (1x) cells. (**c**) Dynamic range of protein abundance in mini pools overlaid with detection in single cells. The inset Venn diagram summarises the overlap between the proteome from the mini-pool and the single-cell datasets, indicating the number of proteins detected exclusively in each set or shared between them. (**d-e**) Enrichment of selected neuronal markers in the top 10% most abundant proteins, comparing phosphorylated tau-negative (left) *versus* positive (right) samples in mini-pool (D) and single-cell (E) datasets. (**f**) Cell-type enrichment of top 100 most abundant proteins detected in the mini-pool dataset of 20x cells showing significant enrichment in superficial excitatory neurons (population named “Ex3e”). (**g**) Tau levels in 20x cells (mini-pool dataset) and single-cells are significantly increased in pTau positive samples. Two-way ANOVA followed by Tukey’s pairwise *post hoc* tests. (**h**) Detection rate (%) of tau peptides plotted along the amino acid sequence of the 2N4R isoform (441 aa), with individual peptides shown as points and smoothed trend lines for pTau-negative (blue) and pTau-positive (yellow) cells across major functional domains. (**i**) Fraction of pTau217-positive cells among the pTau captured cells (AT8-positive) from the mini-pool dataset (left); fraction of pTau231-positive cells among the pTau captured cells (AT8-positive) from the mini-pool dataset (right). (**j**) Representative immunostainings from layer II of BA9 neurons stained with AT8 (green) and pTau217 (magenta). Scale bar: 50µm (left). Quantification of the proportion of AT8 and pTau217 positive cells in 10 AD cases (right). (**k**) Representative immunostainings from layer II (BA9) neurons stained with AT8 (green) and pTau231 (magenta). Scale bar: 50µm (left). Quantification of the proportion of AT8 and pTau231 positive cells in 8 AD cases (right). Paired t-test t(7)=-2.48. (**l**) pTau231 (log₂ intensity) levels in pTau-negative and pTau-positive cells from the mini-pool dataset. Unpaired t-test t_42_=7.34, p=4.72e-9. (**m**) Residuals from a linear mixed-effects model predicting pT231 intensity from total tau abundance for both pTau-negative and pTau-positive cells, with elevated residuals in pTau-positive cells (linear mixed-effect model; β=1.16; p<0.001).

**Table 1.** Case demographics.

| Case # | PMI hours | Age at death | Sex | Clinical Dg | Path | TDP43 limbic | Thal | Braak | CERAD | ABC | Included in SC |
| --- | --- | --- | --- | --- | --- | --- | --- | --- | --- | --- | --- |
| 1 | 31.42 | 63 | M | AD | AD | No | 5 | 6 | 3 | A3B3C3 |  |
| 2 | 74.4 | 63 | M | CBS due to AD | AD | No | 5 | 6 | 3 | A3B3C3 | Yes |
| 3 | 34.2 | 64 | M | AD | AD | No | 5 | 6 | 3 | A3B3C3 |  |
| 4 | 104.55 | 65 | M | AD | AD | No | 5 | 6 | 3 | A3B3C3 | Yes |
| 5 | 35.2 | 68 | M | bvFTD | AD | No | 5 | 6 | 3 | A3B3C3 |  |
| 6 | 73.45 | 68 | M | AD | AD | No | 5 | 6 | 3 | A3B3C3 | Yes |
| 7 | 72 | 69 | M | CBS | AD | No |  | 5 | 3 | A3B3C3 |  |
| 8 | 35.04 | 69 | M | AD | AD | No | 5 | 6 | 3 | A3B3C3 | Yes |
| 9 | 84.45 | 69 | M | PPA | AD | No | 5 | 6 | 3 | A3B3C3 |  |
| 10 | 95.1 | 86 | M | PPA | AD | No | 5 | 5 | 3 | A3B3C3 | Yes |

We identified up to 2,677 proteins in single cells and 5,530 in mini-pool samples, with median proteome depths of 1,698 proteins and 5,060 proteins, respectively (**Fig. 1b**). Proteomes from single-cells were enriched for the most abundant proteins identified in mini-pool samples, with the two datasets sharing more than 95% overlap. (**Fig. 1c**). Disease-relevant proteins, including the microtubule-associated protein tau and p62/SQSTM1, a selective autophagy adaptor protein that binds tau aggregates (Ferrari et al. 2024), were robustly quantified in both single and pooled neuron sets, demonstrating sufficient sensitivity to capture key pathological markers (**Fig. 1c**).

### Atlas-Based Cell-Typing Confirms Upper Layer Neuronal Identity

NFTs preferentially accumulate in excitatory neurons (Fu et al. 2017, 2019; Hyman et al. 1984; Kowall and Beal 1991). Consistent with this, we confirmed the neuronal identity of both pTau-positive and negative cells, evidenced by their enrichment and similar levels of canonical neuronal markers (*i.e.* CAMK2A, SNAP25, SLC17A7, MAP2, PRNP) (**Fig. 1d,e**). In addition, we verified cell-type identity of captured neurons by EWCE (Expression Weighted Celltype Expression), which leverages public single-cell RNA sequencing atlases as reference datasets for cell-type enrichment analysis (Skene and Grant 2016). Results revealed that the 100 most abundant proteins in the mini-pool dataset (85 of which were also among the top 15% most abundant proteins in single-cell dataset), were significantly enriched for excitatory neuronal markers specific to superficial cortical layers (II-III) (**Fig. 1f**). This indicates that our pipeline enables precise cell-type isolation, resulting in a homogeneous dataset.

### Distinct tau phosphorylation states coexist within the same cortical layer

To profile tau in pTau positive *versus* negative neurons, we first compared relative tau protein levels between the two cell populations. Total tau was significantly increased in pTau positive neurons (**Fig. 1g**), consistent with tau accumulation in NFTs. Peptide coverage analysis showed detection of peptides across the full length of the tau protein (isoform 2N4R), with approximately 48% also identified at single-cell level. No preferential enrichment of 3R or 4R tau isoforms was observed, however the N-terminus was more represented in pTau-positive neurons (**Fig. 1h, Extended Fig. 1a**). This pattern may reflect increased solubility and digestibility of the N-terminal region, reduced post-translational modification relative to the C-terminus, or the presence of proteolytically truncated tau species that retain the N-terminus, while lacking portions of the C-terminal region. Fifteen peptides were significantly increased in pTau-positive samples (**Extended Fig. 1b).**

Given that tau is extensively post-translationally modified, including phosphorylation at sites implicated in early NFT formation (Köpke et al. 1993; Kumar et al. 2026; Bancher et al. 1989), we searched MS data using a predicted phospho-library. We identified four high-confidence tau phospho-peptides (site localization probability ≥0.99; **Supplementary Table 1** and **Extended Data Fig. 2a)**, with pT231 being the most abundant (42.25%), followed by pS202 (29.58%), pT217 (26.76%), and pS404 (1.41%). In the mini-pool data, pT217 was detected in ∼80% of pTau-positive and none of the pTau-negative samples, while pT231 was present in all pTau-positive and ∼35% of pTau-negative neurons (**Fig. 1i**). Immunofluorescence confirmed these patterns: neurons positive for pT217 but negative for AT8 (pTau) were not observed, whereas pT231-positive neurons were significantly more frequent than AT8-positive neurons (paired t-test: t₇=2.38, p=0.0419), with approximately 23.98% of neurons showing an AT8⁻/pT231⁺ profile (**Fig. 1j,k**). Quantitatively, pT231 levels in the mini-pool data were significantly elevated in pTau-positive samples (β=54,081, p=4.14e-19, **Fig. 1l**).

To determine whether phosphorylation increases independently of total tau levels, we modelled pT231 as a function of total tau in pTau-negative cells. pT231 levels in pTau-positive neurons exceeded abundance-based expectations by 2.2-fold (β=1.16 log₂ units, p<0.001; **Fig. 1m**), indicating increased site occupancy beyond tau accumulation.

We also identified phosphorylation at S202 and S404 (**Extended Data Fig. 2a**). pS202 is within the AT8 antibody domain (S202/S205). Although the peptide containing S205 was detected, phosphorylation at this site was not observed in our dataset most likely reflecting technical limitations of mass-spectrometry *versus* immunofluorescence or low phosphosite stoichiometry. pS404 detection is consistent with more advanced pathological tau recognized by late-stage tangle antibody PHF1 (pS396/S404) (Augustinack et al. 2002). Overall, pTau-positive neurons show increased tau without isoform-specific enrichment, and within the same brain we identified different distributions of phosphosites, indicating heterogeneity in tau phosphorylation state across neurons.

In addition to tau, several other phospho-proteins were identified (**Supplementary Table 1** and **Extended Data Fig. 2b**). The accumulation of these phospho-peptides is consistent with altered cytoskeletal dynamics (NEFM, ADD1/2. STMN1), modulation of synaptic vesicle machinery (STX1B 14, MYO5A and DNAJC5), engagement of protein quality control systems (HSPB1, HSP90AB1, CRYAB, DNAJC5, TRIM2) and mitochondrial metabolism and degradation (PDHA1 and BCL2L13).

### Proteomic Response To Tau Accumulation

To understand the molecular consequences of tau accumulation, we next analyzed the proteome profile between pTau positive and negative neurons. Low-dimensional representation of the mini-pool proteome in UMAP representation revealed a clear continuum between pTau-positive and pTau-negative neurons, rather than discrete clusters (**Fig. 2a**). Based on this, we reconstructed a pseudotime trajectory using nearest-neighbor graphs (Setty et al. 2019), ordering neurons along a continuous axis such that proteomically similar cells are positioned closer together. This approach not only captures gradual transitions between states that would be missed by binary classifications (*e.g.* pTau+ *versus* pTau-) but also allows us to study at single-cell resolution how molecular changes develop as tau pathology emerges and progresses (**Fig. 2a,b**).

**Figure 2.**
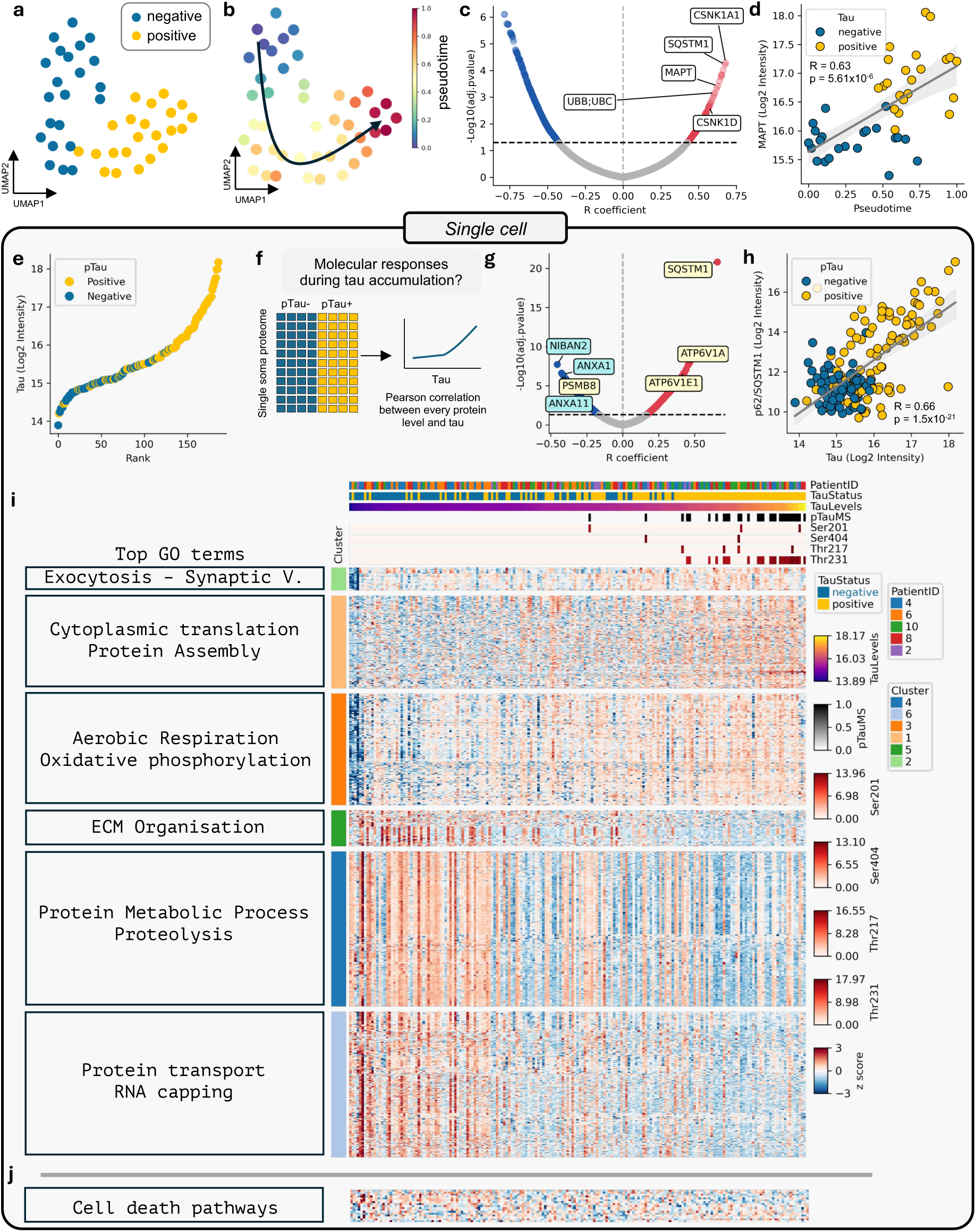
Molecular responses to tau accumulation. (**a**) Low dimensional embedding (UMAP) of positive (yellow) and negative (blue) neuron proteomes. (**b**) Pseudotime-based ordering of negative to positive neurons representing a pathology axis. (**c**) Volcano plot showing the correlation strength (Pearson’s R coefficient) of individual proteins with pseudotime in the mini-pool dataset. Six representative highly correlated proteins were selected among the top candidates and annotated. (**d**) *MAPT* (tau) log₂ intensity plotted against pseudotime in the mini-pool dataset. (**e**) Ranked tau (log₂ intensity) values across the single-cell dataset. (**f**) Schematic illustrating the analytical framework used to investigate molecular responses during progressive tau accumulation, with cells ordered along a continuum from low (blue) to high (yellow) tau levels. (**g**) Volcano plot showing the correlation strength (Pearson’s R coefficient) of individual proteins with tau in the single-cell dataset. Seven highly correlated proteins were selected from the top candidates and annotated, with proteins implicated in cell-death mechanisms shown in blue and those associated with the proteasome highlighted in yellow. (**h**) p62/SQSTM1 (log₂ intensity) plotted against total tau (log₂ intensity) in the single-cell dataset. (**i**) Clustered heatmap of significantly correlated proteins (FDR < 0.05). Cells (columns) are ranked by tau levels. pTau status (including site-specific tau phosphorylation) and patient ID are indicated. Top GO terms are shown for each cluster. (**j**) Heatmap of proteins associated with cell-death pathways in single-cell dataset.

We next examined which proteins were associated with the pseudotime axis (**Fig. 2c; Supplementary Table 2**). Total tau and pT231 displayed strong positive correlations with pseudotime (r=0.63, p=5.61e-6 and r=0.63, p=5.48e-6, respectively; **Fig. 2d, Extended Data Fig. 3a**). Consistent with impaired protein degradation pathways, p62/SQSTM1 and ubiquitin (UBB; UBC) were also positively correlated with pseudotime (ranked #3 and #10, respectively; **Fig. 3c**). Notably, two casein kinase 1 isoforms were among the top 20 most correlated proteins: casein kinase 1α (CSNK1A1; #1) and casein kinase 1δ (CSNK1D; #17). Both isoforms are known to phosphorylate tau at the AT8 (S202/S205) and PHF1 (S396/S404) epitopes and modulate its aggregation (Kuret et al. 1997; Kannanayakal et al. 2006; Li et al. 2004). CSNK1D is a recognized marker of granular vesicular bodies (GVBs), neuron-specific lysosomal structures that arise in response to intracellular protein aggregation and recognized as a neuropathological feature of AD (Köhler 2016), thereby linking its enrichment along pseudotime to the emergence of GVB pathology.

**Figure 3.**
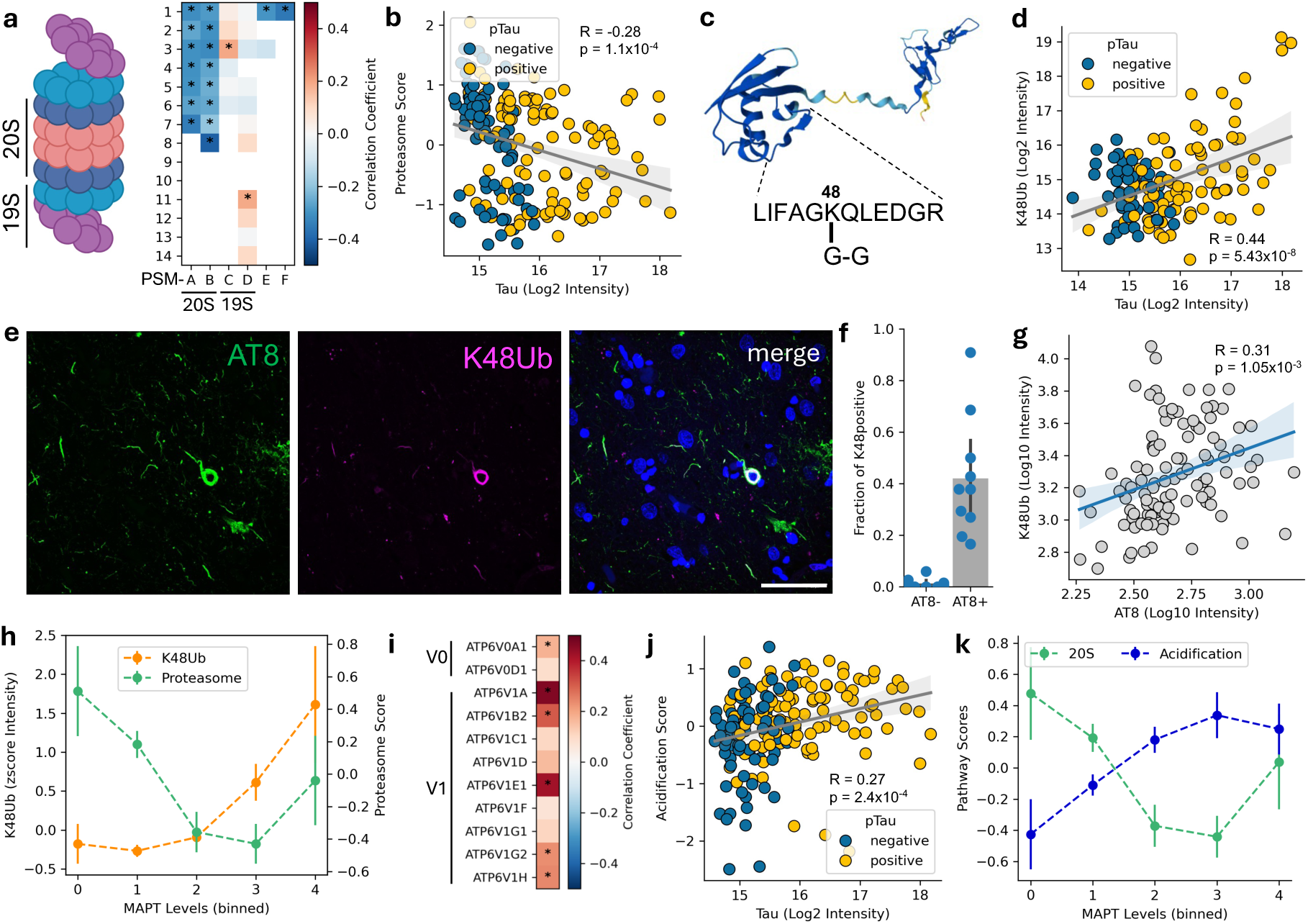
Progressive rewiring of the proteostasis network with tau accumulation. (**a**) Drawing of the proteasome with annotation of the 20S and 19S subunits (left). Heatmap of the individual correlation values for each proteasome subunit identified (right).* indicates proteome-wide (FDR<0.05) statistical significance. (**b**) Scatter plot highlighting the negative correlation between proteasome score and tau levels in single cell proteomics. (**c**) AlphaFold 3D rendering of the ubiquitin protein (RPS27A), the identified peptide, and the location of Gly-Gly (G-G) tag on Lys48. (**d**) Scatter plot highlighting the positive correlation between K48-ubiquitin and tau levels in single cell proteomics. (**e**) Representative immunostainings from layer II (BA9) neurons stained with AT8 (green) and K48 (magenta) and (**f**) quantification showing fraction of K48-positive cells is shown for AT8-negative and AT8-positive cells. (**g**) K48-linked ubiquitin (log₁₀ intensity) plotted against AT8 (log₁₀ intensity). (**h**) Comparison between trends in proteasome score (green) and K48-ubiquitin levels per bin of tau (*MAPT*) levels. (**i**) Heatmap of the individual correlation values for each vATPase subunit identified.* indicates proteome-wide (FDR<0.05) statistical significance. (**j**) Scatter plot highlighting the positive correlation between the acidification machinery score and tau levels in single cell proteomics. (**k**) Comparison between trends in 20S proteasome score (green) and acidification machinery score (blue) per bin of tau (*MAPT*) levels, showing differential responses across tau levels.

Gene ontology analysis of significantly correlated proteins revealed that terms related to extracellular matrix (ECM) organization were among the most negatively enriched (**Extended Data Fig. 3b**). To further resolve these patterns, we performed Gene Ontology analysis on hierarchical clustering of pseudotime-correlated proteins. Cell–matrix adhesion (related to ECM organization) and proteolysis, were negatively correlated with pseudotime, whereas mitochondrial gene expression and regulation of autophagy showed a positive correlation (**Extended Data Fig. 3c-g** and **Supplementary Table 2**).

Given the strong correlation between tau and pseudotime in the mini-pool dataset, together with the close relationship between tau abundance and pTau immunopositivity (**Fig. 2e**), we used tau abundance as a proxy for pseudotime in the single-cell dataset. This approach defined a continuum of neurons spanning from absent or low to high tau levels, enabling the analysis of molecular responses to progressive tau accumulation at a single cell level (**Fig. 2f**). p62/SQSTM1 was the most positively correlated protein (r=0.66, p=1.5e-21; **Fig. 2h**), closely tracking tau levels. This was followed by V1 subunits of the lysosome proton pump (vacuolar ATPases: ATP6V1A and ATP6V1E1). Stress-responsive adaptive protein NIBAN2 was the most negatively correlated protein (r=-0.46, p=1.93e-8). Clustering all 1,062 significant proteins (49.8% of the proteome; **Supplementary Table 3**), followed by gene ontology analysis (**Fig. 2i** and **Extended Data Fig. 4**) revealed functional groups similar to those in the mini-pool analysis, supporting the use of tau abundance as a proxy for pseudotime. Exocytosis, oxidative phosphorylation, cytoplasmic translation and protein assembly were positively correlated with tau levels, whereas extracellular matrix organization, protein metabolism and proteolysis, and protein transport and RNA capping were negatively correlated. Protein transport and translation were detected in the single-cell dataset, highlighting the complementarity of both approaches (**Supplementary Table 4**).

### No detectable activation of canonical cell-death programs in pTau positive neurons

Given the role of NIBAN2 as a pro-survival factor (Chen et al. 2011), we hypothesized that its downregulation could lower the apoptotic threshold, prompting a systematic investigation of cell-death related pathways. Supporting this hypothesis, additional proteins involved in apoptosis regulation also showed negative correlations with tau levels, including Annexin A11, a vesicular trafficking protein implicated in Amyotrophic Lateral Sclerosis (Smith et al. 2017), and Annexin A1, a Ca²⁺-dependent phospholipid-binding protein (**Fig. 2g**). Annexin A1 is a cytosolic protein that can be secreted (de Souza Ferreira et al. 2025), which may explain its apparent decrease in the single-cell proteomics dataset. Together with the absence of functional enrichment for cell-death pathways, these observations prompted us to directly investigate cell-death mechanisms. To obtain an unbiased view of cell-death pathways, we leveraged an AI data analyst (see Methods), which identified candidate proteins associated with multiple cell-death related pathways, including apoptosis, necroptosis, pyroptosis, ferroptosis, and senescence. Only a few proteins (7) were detected in the single-cell dataset, and none showed a clear correlation with tau levels (**Fig. 2j**). To further interrogate this, we also investigated cell-death pathways in the mini-pool data, where 30 cell-death related proteins were quantified (**Extended Data Fig. 5a**). These were compared to a publicly available single-soma transcriptomic dataset comparing AT8 (tangle) positive and AT8 negative neurons (Otero-Garcia et al. 2022). Consistent with previous reports, we observed a modest upregulation of cell-death pathway components at the transcript level (Otero-Garcia et al. 2022). However, we observed a consistent downregulation at the protein level (**Extended Data Fig. 5b**). Together, these findings suggest a molecular adaptation to tangle formation without clear evidence of cell-death induction.

### Rewiring Towards Lysosomal Proteolysis

One of the most negatively correlated proteins with tau levels was PSMB8 (**Fig. 2g**) which is one of the three subunits of the 20S immunoproteasome, previously reported to colocalise with alpha-synuclein aggregates in Parkinson’s Disease (Nguyen et al. 2025; Ugras et al. 2018). Considering the established role of the proteasome in protein homeostasis control and its impairment in tauopathies (Myeku et al. 2016; Collins et al. 2025; Jiang et al. 2025), we performed an extended investigation of proteasome subunits. All subunits of the 20S (*i.e.* catalytic) proteasome negatively correlated with tau levels (**Fig. 3a,b** and **Extended Data Fig. 6a**). We next built a proteasome score by averaging the levels of all significantly correlating proteasome subunits (**Fig. 3b**). In the canonical configuration of the 26S proteasome, the 20S core is flanked by two regulatory 19S subunits (Finley 2009). Two members of the 19S (PSMC3 and PSMD11) displayed a significant positive correlation with tau levels, but most other members remained unchanged (**Fig. 3a**) highlighting a differential effect on 20S and 19S subunits in AD as previously reported (Collins et al. 2025).

To probe for proteasome activity, we reasoned that such a downregulation of the catalytic core components would lead to the accumulation of ubiquitinated substrates. Similar to our phosphoproteomic search, we re-analyzed our data for additional ubiquitin modifications using a predicted K-GG peptide spectral library (*e.g*. Gly-Gly tags), which identified the K48-linked Gly-Gly remnant on ubiquitin with high confidence (encoded by RPS27A; **Fig. 3c**). This peptide is indicative of K48-linked poly-ubiquitin chains, the canonical signal for proteasomal degradation, and it strongly correlates with tau and pTau231 levels (**Fig. 3d**), further supporting impaired proteasome function with increasing tau. We confirmed this observation using immunostaining against K48-ubiquitin and pTau (**Fig. 3e**). We found that K48 staining was mostly confined to AT8-positive cells with a large fraction of AT8-positive cells displaying K48 accumulation and a significant positive correlation between AT8 levels and K48 (**Fig. 3f,g**, r=0.31, p=1.5e-3). Binning cells by tau levels revealed an early decrease of proteasome subunits, followed by elevation of K48-ubiquitin (**Fig. 3h**). Of note, the decreased proteasome score was strongest at bins 0 and 1, which are predominantly composed of pTau-negative cells, suggesting that proteasome function is affected prior to the accumulation of AT8 positive tau.

The ubiquitin-proteasome system and the autophagy-lysosomal pathway are major components of the neuronal proteostasis network (Labbadia and Morimoto 2015). Having established a decline in proteasome capacity with increasing tau, we next examined the response of lysosomal components. Two subunits of the cytosolic regulatory V1 subcomplex of the lysosomal proton pump (ATP6V1A and ATP6V1E1) were amongst the most positively correlated proteins with tau (**Fig. 2g**). Lysosomes derive their degradative functions from their acidic pH (∼4.5), which is critical for the proper activity of luminal hydrolases. An extended investigation of all subunits of the vacuolar-ATPase (V-ATPase) demonstrated that all detected subunits, spanning both V0 and V1 subcomplexes, were positively correlated with tau levels (**Fig 3i** and **Extended Data Fig. 6b**). We built an acidification score by averaging the normalized abundance levels of all detected subunits (**Fig. 3j**), which highlighted an early upregulation of V-ATPase components (**Fig. 3k**), mirroring the decrease in 20S proteasome subunits and revealing an early rewiring of the proteostasis network.

This observed increase of lysosomal acidification machinery contrasted with other studies of lysosomal acidification in AD assessed by bulk tissue proteomics, where a decrease was observed, coupled with a reduction in proteolytic activity (Nixon et al. 2024; Lee 2022; Kim et al. 2023). To investigate whether the discrepancy could be related to cellular resolution, we screened the transcriptomics dataset of tangle-bearing neurons (Otero-Garcia et al. 2022). Confirming our proteomics observations, the single-soma transcriptomics data also showed increased expression of most subunits of the lysosome proton pump (**Extended Data Fig. 6c**). While bulk tissue analyses capture responses across many cell types, our data reveal that individual neurons accumulating tau switch from proteasome-mediated degradation towards lysosomal proteolysis.

### Temporal Ordering Of Molecular Responses To Tau Accumulation

Building on the emergence of AI-based scientific assistants, we leveraged Kosmos, a recently described AI scientist (Mitchener et al. 2025), to perform an unbiased analysis capable of extracting patterns and relationships that may not be apparent through standard methods. Kosmos deploys multiple specialized agents (*e.g.* for literature search or data analysis) that iteratively integrate their outputs into a structured world-model that evolves over iterations (Mitchener et al. 2025).

We provided the single-cell dataset to Kosmos and tasked it with proposing mechanisms underlying tau accumulation and reconstructing a temporal sequence of associated molecular changes (**Fig. 4a**). Kosmos suggested that proteins changing with disease progression (tau accumulation) may not follow simple linear patterns (upregulated or downregulated), but instead show shifts in abundance at specific points. Identifying these “breakpoints” can help estimate when particular molecular changes occur and therefore pathway responses. We iteratively refined the analysis by combining AI suggestions with expert input (“scientist-in-the-loop” framework described in (Mitchener et al. 2025)). For each protein, we fitted models that were either linear or allowed one or two breakpoints, selecting the best fit using Bayesian information criterion. This resulted in the classification of four patterns (**Fig. 4b**): (i) proteins that increase with tau levels (*e.g*. p62/SQSTM1); (ii) proteins that decrease with tau levels (*e.g*. NIBAN2); (iii) proteins that show an early breakpoint, indicative of early responses; and (iv) proteins that show a late breakpoint then decline/recover, indicative of delayed responses. Most proteins followed linear profiles, but ∼250 proteins displayed non-linear dynamics (**Fig. 4c**). Among proteins with a single breakpoint, we observed two main groups, corresponding to early and late changes along the tau accumulation axis (breakpoints at log₂ MS intensity ∼15.2 and ∼16.7, respectively; **Fig. 4d**).

**Figure 4.**
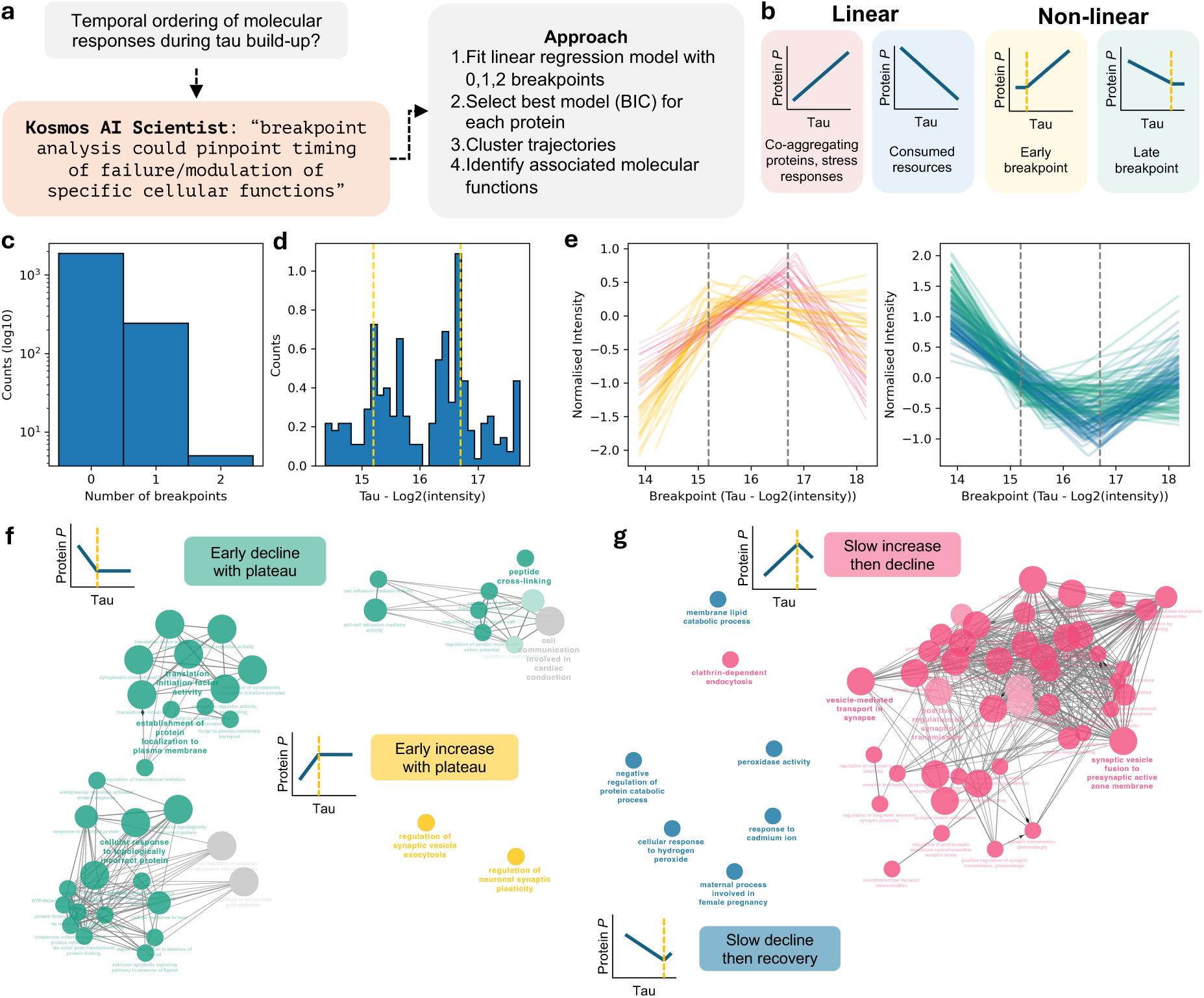
AI-inspired breakpoint analysis allows temporal ordering of pathological events. (**a**) Graphical summary of Kosmos output and the approach for temporal ordering. (**b**) Possible scenarios of linear and non-linear correlations between tau and individual proteins. The identification of a possible breakpoint allows ordering of the events. (**c**) Distribution of number of breakpoints per protein. (**d**) Distribution of breakpoint positions (log₂ tau intensity) for proteins with one breakpoint. The yellow lines highlight the local maxima and define early and late breakpoints. (**e**) Profiles of proteins with one breakpoint showing upregulation (left) or downregulation (right), stratified by early and late breakpoints. (**f,g**) Functional annotation of the different profiles for early (f) and late (g) breakpoints. Each node represents a gene ontology term, scaled based on the number of members. Edges highlight the number of members shared between nodes. Nodes are coloured based on the different profiles from (e).

Gene Ontology analysis on the trajectories revealed four main protein modules. Two modules showed early changes: proteins that increased then plateaued (yellow) and those that decreased then stabilized (green) (**Fig. 4e,f** and **Supplementary Table 5**), consistent with early adaptive or vulnerability responses to rising tau. The remaining two modules encompassed proteins showing slow changes followed by decline (pink) or recovery (blue) (**Fig. 4e, g** and **Supplementary Table 5**).

The yellow and pink modules were enriched for synaptic proteins but showed different temporal dynamics and functional roles. Specifically, the early rise of the yellow module (**Fig. 4e, f**) was associated with proteins involved in synaptic regulation and signaling, including GRIN1 (NMDA receptor subunit), PRKCG (activity-dependent kinase signaling), RAB3A (regulation of vesicle release probability), and CACNA2D1 (calcium handling). Together, these changes suggest an early compensatory phase characterized by synaptic plasticity, rewiring of the synaptic system, in which neurons actively adjust synaptic connectivity to maintain function under tau-induced stress. This is supported by data showing a silencing and rewiring of tangle-bearing neurons (Zwang et al. 2025). The subsequent plateau suggests that this adaptive capacity reaches a functional limit despite continued tau accumulation.

The pink module (**Fig. 4e,g**) involved core components of the presynaptic vesicle cycle (*e.g.* SNAP25, STX1A/B, STXBP1, SYP, SV2A, SYN1, SYT1) and endocytic machinery (DNM1/3, CLTA), indicating increased demand on vesicle recycling. This suggests a slow attempt to sustain neurotransmission under escalating stress. However, at the late breakpoint, the pink module declined, consistent with a failure of synaptic machinery. The enrichment of presynaptic proteins is in line with reports of tau accumulation at presynaptic terminals and subsequent synapse loss (Zhou et al. 2017; Kiani 2023; Fertan et al. 2026).

The green module (**Fig. 4e,f**) reflected a stressed, proteostasis-activated state in which translation, degradation, and trafficking pathways were affected as a consequence of early tau accumulation. The decline in core translation factors (EIF2S1, EIF3A/B/C, EIF4A2, EEF2) suggests suppression of global protein synthesis, while the concurrent decrease in chaperones and proteasome components (HSPA1A/B, HSPA5, PSMA6, PSMB3) points to a diminished ability to refold or degrade misfolded proteins. This pattern is consistent with activation of the integrated stress response, whereby cells suppress global protein synthesis to prevent further overload of the proteostasis machinery, a process previously linked to tau pathology (Bond et al. 2020; Pitera et al. 2021).

The blue module (**Fig. 4e,g**) showed early suppression followed by recovery. In early stages, proteins involved in proteostasis (PSMA/PSMB, UBB/UBC, USP7), lysosomal degradation (TPP1, ASAH1), redox regulation (CAT, TXN), and ER/secretory function (SSR1/4, DDOST) were downregulated, suggesting impaired cellular quality control and metabolic homeostasis, potentially as an adaptive response to rising tau. Beyond the late breakpoint, these pathways were broadly upregulated, suggesting a late response involving protein degradation, oxidative defense, and protein folding/trafficking, alongside cytoskeletal and inflammatory changes. These dynamics are consistent with prior reports showing early impairment of proteostasis, lysosomal, and redox pathways in tauopathy, followed by late compensatory stress activation.

## Discussion

Understanding how neurons respond to tau accumulation remains a central challenge in Alzheimer’s disease research. Most proteomic studies of AD have relied on bulk tissue measurements, averaging signals across thousands of cells with heterogeneous pathological states. In this study, we define a quantitative proteomic trajectory of neuronal response to tau pathology between neurons in proximal regions within the same Alzheimer’s disease brain, revealing that neurons respond differently depending on the degree of tau accumulation. Rather than separating into pTau-positive and pTau-negative classes, neurons organize along a continuum of molecular states that correlate with tau abundance, revealing a staged response in which distinct pathways are engaged at progressively higher levels of tau burden. Within this continuum, some pathways show consistent directional changes, being either upregulated or downregulated as tau accumulates, including ECM organization, oxidative phosphorylation, proteolysis, and RNA capping. In contrast, other pathways exhibit staged, non-linear dynamics that become apparent when modelling protein trajectories using breakpoint analysis, highlighting subtle functional changes. For example, temporal dynamics across the identified modules indicate that synaptic dysfunction in tauopathy is not a single event but a staged process associated with increasing tau burden and distinct molecular signatures. Together with the progressive rewiring of proteostasis pathways, these findings support a model in which tau pathology drives sequential phases of adaptation, compensation, and eventual functional collapse.

An interesting finding was the coordinated remodeling of the neuronal proteostasis network. All detected subunits of the catalytic 20S proteasome core negatively correlated with tau levels, while ubiquitin conjugates indicative of K48-linked polyubiquitination accumulated with increasing tau. These observations indicate impaired proteasomal degradation in neurons with rising tau burden. In contrast, the lysosomal acidification machinery, including multiple V-ATPase subunits, showed the opposite response, increasing early along this trajectory. This reciprocal regulation points to a shift from proteasome-mediated degradation toward lysosomal proteolysis as tau accumulates. Such a switch may reflect a compensatory response to declining proteasome capacity, consistent with prior reports implicating lysosomal pathways in the clearance of aggregated proteins.

An important implication of this work is that not all pTau-negative neurons in the AD brain are unaffected, but they instead occupy early positions along the molecular trajectory of disease progression. These low-tau/pTau negative neurons were enriched for proteins involved in ECM, proteolysis, and protein transport/RNA capping, indicating that substantial molecular remodeling precedes overt pTau accumulation. These pathways could define an early vulnerability state that could lead to initiation of the pathological cascade, or an early response to some form of pathological tau that is not captured by AT8 immunopositivity.

This study identifies several advancements. First, it demonstrates the feasibility of performing exploratory proteomics on archival FFPE human AD brain tissue at single-neuron resolution, allowing for the study of molecular pathology retrospectively in clinically and neuropathologically characterized cohorts. Second, the combination of single-cell and mini-pool proteomics provides complementary insight: single-cell data captures neuron-to-neuron heterogeneity and enables trajectory-based analyses, whereas mini-pools provide greater proteome coverage including tau proteoform information. To reinforce this, key molecular trends were consistent across both modalities, including the strong positive correlation of p62/SQSTM1 with tau levels and the inverse relationship between tau abundance and proteasome subunits, supporting the robustness of the proteostatic remodeling data. Third, we directly identified phosphorylated proteins whose modification states appear to regulate pathways highlighted by the cluster analysis; this underscores a key advantage of proteomics over transcriptomic approaches in capturing post-translational functional regulations. In addition, the direct detection of disease-relevant phospho-tau peptides in such low input material, including at the level of individual neurons, highlights the sensitivity of the workflow. Fourth, the integration of an AI-guided analytical framework offered an unbiased perspective on the data, and highlighted non-linear temporal responses that may not have emerged from standard analysis alone. Finally, by modelling neurons along a tau continuum rather than forcing them into binary pTau-positive *versus* pTau-negative categories, we were able to resolve subtle, progressive molecular changes that would likely be lost in a dichotomous comparison.

This continuum-based approach also provides a conceptual advance over the traditional framework of inferring disease progression by comparing individuals at different Braak NFT stages. Braak staging remains invaluable for identifying anatomical spread of pathology across the brain, but it does not capture the burden, or full heterogeneity, of pathological state among neurons within the same region of the same individual. By focusing on neurons from the same Braak stage and even from the same local microenvironment, our analysis minimizes confounds that arise when comparing controls with affected cases, or early-stage with late-stage cases, and show that even within a single Braak stage, a substantial biological heterogeneity exists within a region that is already classified as pathologically involved. This intra-regional heterogeneity has important therapeutic implications. If neurons at late Braak stages still occupy early positions along the tau-response trajectory, then intervention may remain beneficial even in advanced disease. Such cells may be especially amenable to therapies aimed at restoring proteostasis, improving mitochondria metabolism, or modulating cell-matrix signaling pathways. Both oxidative phosphorylation and extracellular matrix pathways have been linked to neuronal resilience in a previous study (Lin et al. 2025), further supporting their potential as targets for therapeutic intervention. This view argues against an overly deterministic interpretation that late Braak stage indicates late-stage pathology and supports the possibility that some neuronal populations remain biologically rescuable even after disease has become widespread.

The temporal response model may not necessarily be interpreted as a simple linear sequence through which every neuron passes in the same order. An alternative interpretation is that different neurons may preferentially engage different response programs. Some neurons may mount an early-response program, including synaptic rewiring and adaptation, and ultimately prove more resilient to tau pathology. For example, synaptic plasticity and calcium homeostasis have been shown to be upregulated in early-stage resilient neurons (Dharshini et al. 2026). Others may enter later-response states reflecting a more vulnerable trajectory. Under this view, the four temporal response patterns do not necessarily define obligatory sequential steps but may instead reflect partially distinct neuronal fates. This possibility could also help explain why we detect extensive molecular remodeling but little clear evidence of overt cell-death pathway activation: if the most vulnerable neurons are underrepresented or already lost, and if the sampled neurons disproportionately capture surviving or adapting cells, then cell death signatures may be obscured.

The relative absence of cell-death pathway activation aligns with emerging literature suggesting that tangle-bearing neurons show reduced risk of cell death (Zwang et al. 2024; de Calignon et al. 2009; Zwang et al. 2023). Our data indicate substantial remodeling of proteostasis, synaptic pathways, redox systems, and trafficking pathways, yet little evidence of canonical apoptosis, necroptosis, pyroptosis, ferroptosis, or senescence at the protein level. Of note, we see elevated levels of GVB-related proteins. GVBs are thought to accumulate in neurons that are resilient to tauopathy-related death (Balusu and De Strooper 2024), where they act as a protective checkpoint, sequestering the activated necrosome within lysosomal structures thereby delaying execution by necroptotic cell death (Balusu and De Strooper 2024). Our observations are consistent with the idea that neurons undergo prolonged adaptation to tau aggregation rather than immediate execution of a cell-death program. Nonetheless, we cannot exclude the possibility that neurons undergoing terminal degeneration were not captured by our analysis, either because their protein content fell below the detection threshold of our methodology, because the region chosen (Layer II dlPFC) is affected later in the disease, or because AT8 immunolabelling and the presence of a nucleus may preferentially identify neurons prior to overt degeneration. Another possibility is that protein abundance alone may not fully capture the molecular state associated with neuronal degeneration, as critical regulatory mechanisms may instead be mediated by post-translational modifications that were not comprehensively detected in our ultra-low input dataset. However, the large number of neurons analyzed should mitigate this potential sampling bias.

Overall, our findings support a model in which tau pathology induces a spectrum of neuronal states that can be resolved within a single diseased brain. Our study reveals that tangle-free neurons can display molecular changes associated with the trajectory of tangle formation, with an early rewiring of the proteostasis network preceding the collapse of synaptic programs as tau accumulates. This molecular timeline opens avenues for targeting early-affected pathways before irreversible neuronal damage occurs.

## Materials and Methods

### Cases

All selected cases were obtained from the Queen Square Brain Bank (QSBB) for neurological disorders (University College London, London, UK). Ethical approval was obtained from the National Hospital for Neurology and Neurosurgery Local Research Ethics committee and the tissue request committee at QSBB in accordance with the human tissue authority’s code of practice and standards under licence number UCLMTA 06-23. Cases met the current diagnostic criteria AD ((Braak and Braak 1991; Thal et al. 2002; Mirra et al. 1991; McKhann et al. 2011; Montine et al. 2012), all reaching a final ABC score of A3B3C3. Demographics and clinical data for all cases used in this study are listed in **Table 1**.

### Immunohistochemistry

Prior to laser microdissection, phosphorylated tau pathology was visualized using antibody AT8, which recognizes phosphorylated tau at S202/T205 and is widely used for neuropathological assessment. Neurons that were negative for tau S202/T205 in cortical layer II were identified with cresyl violet (Nissl) counterstaining, which enabled clear delineation of cortical lamination in the BA9 region of the frontal cortex. Immunohistochemistry was performed as follows. Formalin-fixed, paraffin-embedded (FFPE) frontal cortex (BA9) was sectioned at 7 μm onto UV-treated PPS metal frame slides (Leica). Sections were dried at 40°C and further incubated overnight at 60°C to ensure adhesion. After deparaffinization in xylene and rehydration through graded alcohols, endogenous peroxidase activity was quenched with methanol containing 0.3% H₂O₂. Antigen retrieval was performed in 0.1 M citrate buffer (pH 6.0) for 10 minutes in a microwave, followed by blocking of non-specific binding with 10% milk in PBS. Sections were incubated overnight with a biotinylated AT8 antibody (1:500; Thermo Fisher Scientific) to detect phosphorylated tau, and the signal was amplified using ABC-HRP (Vector Labs) for 30 minutes. Visualization was achieved with 3,3′-diaminobenzidine (DAB) activated by H₂O₂. Slides were counterstained with 0.1% cresyl violet acetate (Sigma), air-dried, and immediately prepared for laser microdissection.

### Laser microdissection

Stained AT8 positive and AT8-negative/cresyl violet-positive neurons were captured using a Leica LMD7 system operated with Leica Laser Microdissection software (v8.3.0.08259). Tissue was dissected using 20x objective and laser settings were as follows: power 56, aperture 1, speed 15, middle pulse count 1, final pulse −1, head current 37–45%, pulse frequency 801, and offset 101. Single-neurons or pools of 20 neurons were collected into a low-binding 384-well plate (Eppendorf) configured over the ‘universal holder’ function. Isolated neurons were mature tangles; pre-tangles and ghost tangles were avoided based on morphology.

### Sample preparation

To ensure settling of dissected contours at the bottom of the plate, wells were washed with 20 µl of 100% LC-MS grade acetonitrile (ACN) and dried in a SpeedVac for 20 min at 45°C. For tissue lysis, 2 µl of buffer (0.1% DDM, 5 mM TCEP, 20 mM CAA and 100 mM TEAB) were added to each well and incubated at 95°C for 60 min. Samples were sequentially digested by adding 1 µl LysC (4 ng/µl in 0.1 M TEAB and 30% LC-MS grade ACN) for 4 h followed by overnight digestion after adding 1 µl trypsin (6 ng/µl in 0.1 M TEAB and 10% LC-MS grade ACN) in a 384-well thermal cycler (BioRad S1000 ThermalCycler) at 37°C. All buffers were dispensed using MANTIS liquid dispenser (Formulatrix). Samples were dried in a SpeedVac for 30 min at 60°C and stored at −20°C until further processed. Before measurement Evotips (Evosep) were prepared and loaded following manufactureŕs instructions. To transfer samples from the 384-well plate into individual Evotips, pipette tips were pre-coated with 0.015% DDM by aspiring and dispensing the solution several times.

### LC-MS/MS analysis

Data acquisition was performed on a timsTOF Ultra 2 mass spectrometer (Bruker Daltonics) coupled to an Evosep One liquid chromatography system (Evosep). Peptides were separated using the pre-programmed gradient for 40 samples per day (SPD) on a 15 cm column with an inner diameter of 75 µm and 1.7 µm C18 beads (Aurora Elite, IonOpticks) maintained at 50°C. The mobile phases comprised 0.1% FA in LC-MS grade water and 0.1% FA in LC MS grade ACN. All samples were acquired with a ramp time of 100 ms in dia-PASEF mode using the manufactureŕs default dia-PASEF windows scheme. In brief, eight dia-PASEF windows with three IM steps covered a m/z range of 400-1000 and an ion mobility range from 0.64-1.37 Vs/cm^2^. The system was operated in high sensitivity mode and ion charge control 2.0 was enabled for a subset of samples. All other parameters were kept as default.

### Raw data analysis

diaPASEF raw file processing was carried out using DIA-NN (version 2.0). The files were searched against an in-silico spectral library generated with a human reference proteome FASTA file (UP000005640_9606) supplemented with common contaminants (Frankenfield et al. 2022). For Tau, we used the isoform Tau-F (P10636-8) corresponding to the brain specific 2N4R isoform. The DIA-NN search included following settings: Protease = ‘Trypsin/P’, Missed cleavages = 2, Maximum number of variable modifications = 1, Mass accuracy = 15, MS1 accuracy = 15, Scan window = 0, Precursor FDR (%) = 1, Scoring = ‘Generic’, Proteotypicity = ‘Genes’, Machine learning = ‘NNs (cross-validated)’, Cross-run normalization = ‘RT-dependent’, Library generation = ‘Smart profiling’, Speed and RAM usage = ‘Optimal results’. Following settings were enabled: N-term M excision, C carbamidomethylation, Ox(M), MBR, Protein inference.

### Raw data analysis for Phosphorylation-/Ubiquitination-sites

For the search of phosphorylated or ubiquitinylated peptides an in-silico digested library was generated in DIA-NN (version 2.0) using a proteome FASTA file (UP000005640_9606). For Tau, we used the isoform Tau-F (P10636-8) corresponding to the brain specific 2N4R isoform of 441 aa. Following settings were used for library generation and subsequent diaPASEF raw file processing: Protease = ‘Trypsin/P’, Missed cleavages = 1, Maximum number of variable modifications = 3, Mass accuracy = 0, MS1 accuracy = 0, Scan window = 0, Precursor FDR (%) = 1, Scoring = ‘Peptidoforms’, Proteotypicity = ‘Genes’, Machine learning = ‘NNs (cross-validated)’, Cross-run normalization = ‘RT-dependent’, Library generation = ‘IDs, RT & IM profiling’, Speed and RAM usage = ‘Optimal results’. Following settings were enabled: N-term M excision, C carbamidomethylation, MBR, Protein inference, XICs and Phospho (for the phospho search) or K-GG (for the ubiquitin/K-GG search). First, all mini-pool samples were processed alone to generate a refined predicted library. Subsequently, all samples including single-cell proteomes were jointly searched using this refined library including either phosphorylation or ubiquitination as variable modification.

### Raw data analysis using diaTracer

Data was searched using diaTracer (version 1.4.9) (Li et al. 2025) via FragPipe (version 23.1). Proteomic workflow ‘DIA_SpecLib_Quant_Phospho_diaPASEF’ was selected. Following settings were used for diaPASEF Spectrum Deconvolution: Delta Apex IM = 0.01, Delta Apex RT = 3, RF max = 500, Corr Threshold = 0.3. ‘Mass Defect Filter’ was not enabled. Remaining settings were kept as default. All samples were searched.

### Immunofluorescence

FFPE frontal cortex (BA9) was sectioned at 7 μm onto positively charged slides, dried at 40 °C, and incubated overnight at 60 °C. Sections were deparaffinized in xylene, rehydrated through graded ethanol, and subjected to antigen retrieval in 0.1 M citrate buffer (pH 6.0) using a pressure cooker. After PBS washes, non-specific binding was blocked with 10% milk in PBS-T. Sections were incubated overnight at 4 °C with primary antibodies against phosphorylated tau (pT231 and pT217; both from Abcam at 1:500), followed by fluorophore-conjugated secondary antibodies for 1 hour at room temperature in the dark. Nuclei were counterstained (*e.g.*, DAPI), and slides were mounted with antifade medium prior to fluorescence imaging.

### Bioinformatic data analysis

Unless otherwise stated, all computational analyses were performed in Python (v3.12.10) using the scientific Python stack (NumPy v.2.2.5, SciPy v.1.15.2, pandas v2.2.3, matplotlib v.3.10.1, seaborn v.0.13.2) using Jupyter notebooks (Harris et al. 2020; Virtanen et al. 2020; McKinney 2010; Waskom 2021; Thomas et al. 2016; Hunter 2007). All analysis code is available GitHub: https://github.com/MathieuBo/Tau_scProteomics. Proteome datasets were pre-processed using Perseus (version 1.6.15.0) (Tyanova et al. 2016). Intensity values for all samples were log2-transformed. Contaminant proteins (identified by the “Cont_” prefix in UniProt accession) were excluded. The tau protein (UniProt P10636) was re-annotated to correspond to the brain-specific 2N4R isoform (P10636-8). For the mini-pool dataset, proteins were retained when identified in at least 70% of samples within either the pTau-positive or pTau-negative group. For the single-cell dataset, proteins with an identification rate of at least 30% in either group were kept. Remaining missing values were imputed using a low-abundance normal distribution per sample (width=0.3, downshift=1.8). Data was stored and processed as AnnData objects (v.0.11.4) (Virshup et al. 2024). Samples were classified as pTau-positive or pTau-negative based on AT8 immunoreactivity at LCM capture. For single-cell data, samples with less than 1,000 proteins were filtered out. In addition, samples with tau abundance below the 5th percentile were excluded to remove low-quality captures. Intensity values of phosphorylated peptides were extracted from the report phosphosite_99 file. For ubiquitinylated peptides, data from the report.parquet file were filtered as follows: Lib.Q.Value < 0.5, Global.Q.Value < 0.5, Lib.Peptidoform.Q.Value < 0.5, Global.Peptidoform.Q.Value < 0.5, Q.Value < 0.01, PEP <= 0.05, Peptidoform.Q.Value < 0.01, PTM.Site.Confidence > 0.9. Only peptides passing the filter criteria were retained. One K48-ubiquitin outlier (maximum intensity value) was set to missing before analysis.

#### Cell-typing analysis

Markers shown in Fig. 1d,e were obtained from the Human Protein Atlas (https://www.proteinatlas.org/). Cell-type enrichment of 100 most abundant proteins (as determined by ranked log2 intensity) in mini-pools was assessed using the Expression Weighted Cell-type Enrichment (EWCE) package (bootstrap n = 50,000; Benjamini–Hochberg FDR) (Skene and Grant 2016) against the Blue Lake et al. 2018 human frontal cortex reference(Lake et al. 2018).

#### Assessment of hyperphosphorylation

To test whether pT231 phosphorylation exceeds what would be expected from tau abundance alone, we fitted a linear mixed-effects model predicting pT231 intensity from total tau abundance (log2 tau intensity) in pTau-negative (AT8−) mini-pool samples only, with donor as a random effect (statsmodels.formula.mixedlm, formula: “pT231 ∼ tau”, groups=PatientID). This model captures the baseline relationship between tau abundance and pT231 phosphorylation in cells without AT8-detectable pathology. Residuals were then computed for all cells (both pTau-positive and pTau-negative). Residuals above zero in pTau-positive cells indicate phosphorylation exceeding abundance-based expectations. The analysis included n=30 mini-pools following filtering for pT231 detection by mass spectrometry.

#### Construction of tau-associated trajectory

In mini-pools, principal component analysis was performed on the full mini-pool proteome using Scanpy (sc.tl.pca). A nearest-neighbour graph was constructed with k=10 neighbors (sc.pp.neighbors) and visualized using UMAP (sc.tl.umap). Diffusion maps were computed using palantir (v1.4.1) with n_components=5 and knn=5 (Setty et al. 2019). Pseudotime was inferred using palantir.core.run_palantir with the most pTau-negative sample as the early cell (cell index ‘15’), four terminal states (indices ‘40’, ‘13’, ‘41’, ‘5’), num_waypoints=100, and knn=5. In single-cell data, tau protein abundance (log2 MAPT intensity) was used as a proxy for pseudotime ordering.

#### Correlation analysis

For each protein, Pearson correlation coefficients were computed against pseudotime (mini-pools) or tau abundance (single cells) across all samples. P-values were corrected for multiple testing using the Benjamini–Hochberg procedure (scipy.stats.false_discovery_control, method=’bh’). Proteins with FDR < 0.05 were considered significantly correlated.

#### Hierarchical clustering and heatmap generation

Significantly correlated proteins were z-score normalised per protein across all samples. Euclidean pairwise distances were computed (scipy.spatial.distance.pdist) and hierarchical clustering was performed using Ward’s method (scipy.cluster.hierarchy.linkage). For the mini-pool dataset, proteins were split into k=4 clusters (scipy.cluster.hierarchy.fcluster, criterion=’maxclust’); for the single-cell dataset, k=6 clusters were used. Heatmaps were generated with PyComplexHeatmap (v.1.8.2) (Ding et al. 2023). Z-score color scale was clipped at ±3.

#### Pathway activity scores

Pathway scores for proteasome subunits, V-ATPase subunits, and other functional groups were defined as the mean of per-protein z-scores (scipy.stats.zscore, computed per protein across all cells) across pathway members. Specifically: the proteasome score comprised PSMA1–7, PSMB1–8, PSME1, and PSMF1; the 20S proteasome score comprised PSMA1–7 and PSMB1–8; the 19S proteasome score comprised detected PSMC and PSMD subunits; and the acidification machinery score comprised ATP6V1G1, ATP6V1G2, ATP6V1B2, ATP6V1C1, ATP6V1E1, ATP6V1A, ATP6V0D1, ATP6V1F, ATP6V0A1, ATP6V1H, and ATP6V1D. Trends along the tau axis, as shown in Fig. 3H,K, were visualized by binning cells into 5 equal-width bins of tau log2 intensity and plotting mean ± SEM of each score per bin.

#### Gene ontology enrichment

Gene Ontology enrichment analysis was performed using GSEApy (v.1.1.11) (Fang et al. 2023) against the GO_Biological_Process_2025 database for each protein cluster.

#### Cell-death pathway analysis

Candidate cell-death proteins were curated from literature by Edison Analysis AI agent spanning five pathways: apoptosis (30 proteins including BAX, BAK1, BCL2, caspases 3/6/7/8/9, CYCS, APAF1, TP53, FAS, FADD, RIPK1), ferroptosis (21 proteins including GPX4, ACSL4, SLC7A11, TFRC, FTH1, FTL), senescence (28 proteins including CDKN1A/2A, TP53, RB1, GLB1, LMNB1, HMGB1), necroptosis (18 proteins including RIPK1/3, MLKL, ZBP1), and pyroptosis (17 proteins including GSDMD/E, CASP1/4, NLRP1/3). Detected cell-death proteins in the single-cell dataset (n=7) and mini-pool dataset (n=30) were visualized as heatmaps ordered by tau abundance.

#### Transcriptomic comparison

To compare proteomic and transcriptomic responses to tau pathology, we re-analyzed the publicly available single-soma RNA-sequencing dataset from Otero-Garcia et al. (Otero-Garcia et al. 2022). Excitatory neurons were subset from the published AnnData object and stratified by SORT status (AT8-positive versus MAP2-only). Cell-death, proteasome, and V-ATPase subunit expression was compared between AT8+ and AT8− excitatory neurons in the Ex01_CUX2-LAMP5 (L2-L3) subtype to match the cortical layer and cell type profiled in our proteomic dataset.

#### AI-assisted hypothesis generation

The mini-pool proteomic dataset was provided to Kosmos (Mitchener et al. 2025), an agentic AI system that deploys specialized agents for literature search and data analysis, which iteratively builds structured world-models. Kosmos was tasked with proposing mechanisms underlying tau accumulation and reconstructing a temporal sequence of associated molecular changes. The system was not provided with a prescribed analytical approach. Over iterative cycles, combining AI suggestions with expert evaluation (“scientist-in-the-loop”), the system converged on the hypothesis that protein–tau relationships may contain discrete inflection points rather than following monotonic trends. This hypothesis was independently validated and operationalized using standard piecewise regression methods (see below). The exact prompt submitted to Kosmos was:

##### Research Objective

This is a proteomic dataset of mini pools of 10 neurons from Alzheimer’s disease cases. Investigate proteome differences between cells with positive and negative tau status. You can use the precomputed pseudotime to order samples. Your task is to propose mechanisms contributing to the accumulation of tau and the link with cellular dysfunction, death, or survival. Propose a sequence of events leading to tau accumulation.

##### Data Description

In the obs dataframe, you will find some metadata about samples (age at death, pseudotime), and the tau status (positive/negative). Use robust statistical approaches accounting for the possible cofactors. Data are already log2 transformed.

#### Breakpoint analysis

To identify non-linear temporal responses during tau accumulation, each protein’s abundance profile along the tau intensity axis (single cells) was modelled with 0, 1, or 2 breakpoints using piecewise linear regression (piecewise-regression package, v1.5.0) (Pilgrim 2021). Prior to fitting, protein intensities were z-score normalized. Outliers beyond ±3 MAD from the median were removed (Winsorisation). For each protein, piecewise linear models with 0, 1, or 2 breakpoints were fitted with n_boot=1,000 bootstrap iterations. The optimal number of breakpoints was selected per protein by minimizing the Bayesian information criterion (BIC), penalizing model complexity. Model parameters extracted included breakpoint positions and slopes before/after each breakpoint (alpha1, alpha2). For visualization, proteins with exactly 1 breakpoint were selected, and those with breakpoint positions between 0.20 and 0.80 of the tau log2 intensity range were retained to avoid edge effects. Proteins were stratified into four trajectory modules based on the direction of slope change (increasing vs. decreasing) and breakpoint timing (early vs. late, defined by the bimodal distribution of breakpoint positions). Gene Ontology enrichment was performed on each trajectory module as described above.

#### Functional network analysis of breakpoint-derived modules

Proteins from the two early breakpoint modules (early up/plateau and early down/plateau) were jointly submitted as two clusters to ClueGO (v2.5.10) (Bindea *et al*. 2009) within Cytoscape (v.3.10.4) (Shannon *et al*. 2003) for pathway enrichment and network visualization; the two late breakpoint modules were similarly submitted together as a separate two-cluster analysis. The ontology source was GO Biological Process (EBI-UniProt-GOA, 25.05.2022 release; 18,085 reference genes; Homo sapiens). Statistical significance was assessed using a two-sided hypergeometric test with Bonferroni step-down correction (p < 0.05), with GO levels restricted to 3–8 and a minimum of 3 genes and 4% coverage per term. Functionally related terms were grouped by kappa score connectivity (threshold = 0.4) and the leading term per group was selected by smallest p-value. For the early-module analysis, 87 genes were submitted (62 + 25), of which 85 (97.7%) were annotated, yielding 39 representative terms in 7 functional groups. For the late-module analysis, 99 genes were submitted (66 + 33), of which 97 (98.0%) were annotated, yielding 53 representative terms in 11 functional groups. Resulting networks were visualized as functionally grouped nodes–edge graphs in Cytoscape (**Fig. 4f,g**).

## Data availability

The mass spectrometry proteomics data have been deposited to the ProteomeXchange Consortium via the PRIDE partner repository (Perez-Riverol et al. 2025) with the dataset identifier PXD076602.

## Funding

Work undertaken by M.B., M.F, K.E.D, S.P, L. S. D, H.D is supported by the UK Dementia Research Institute through UK DRI Ltd, principally funded by the Medical Research Council, K.E.D, M.F and M.B are supported by funding from the Cure Alzheimer’s Fund; K.E.D is supported by the National Institute of Health (NIH), grant award AG063521. M.F. is also funded by Alzheimer’s Research UK (grant award ARUK-RF2023B-015) and Alzheimer’s Association (grant award 24AARF-1244111, which also funds H.D.). M.B. is also funded by a UKRI Future Leaders Fellowship (APP44655). L.K. and F.C. acknowledge funding support by the Federal Ministry of Education and Research (BMBF), as part of the National Research Initiatives for Mass Spectrometry in Systems Medicine (grant agreement No. 161L0222) and funding from the European Research Council (ERC) under the European Union’s Horizon 2020 research and innovation program (grant agreement No. 101115681).

The funders had no role in the study, data collection and analysis, decision to publish or preparation of the manuscript.

We thank our colleagues at the Max Delbrück Center (MDC) for their support and feedback on the manuscript as well as the MDC technology platform ‘Proteomics’. We also thank Janett König for laboratory support performing proteomics sample preparation.

We thank colleagues Bart De Strooper, Hanna Hörnberg, Melissa Birol, Natura Myeku, and Katja Simon for critical feedback on the manuscript.

Tissue was supplied by the UCL Queen Square Brain Bank for neurological disorders.

## Author contributions

Conceptualization: M.F., M.B., L.K., F.C., K.E.D.

Methodology: M.F., M.B., L.K., F.C., K.E.D, S.P., H.D., L.S.D., R.N.

Funding acquisition: M.F., M.B., L.K., F.C., K.E.D.

Writing – original draft: M.F., M.B., L.K., F.C., K.E.D.

Writing – review & editing: M.F., M.B., L.K., F.C., K.E.D., Z.J., L.M., A.Y.

## Competing interests

L.M. and A.Y are employees of Edison Scientific.

---

**Extended Data Figure 1.**
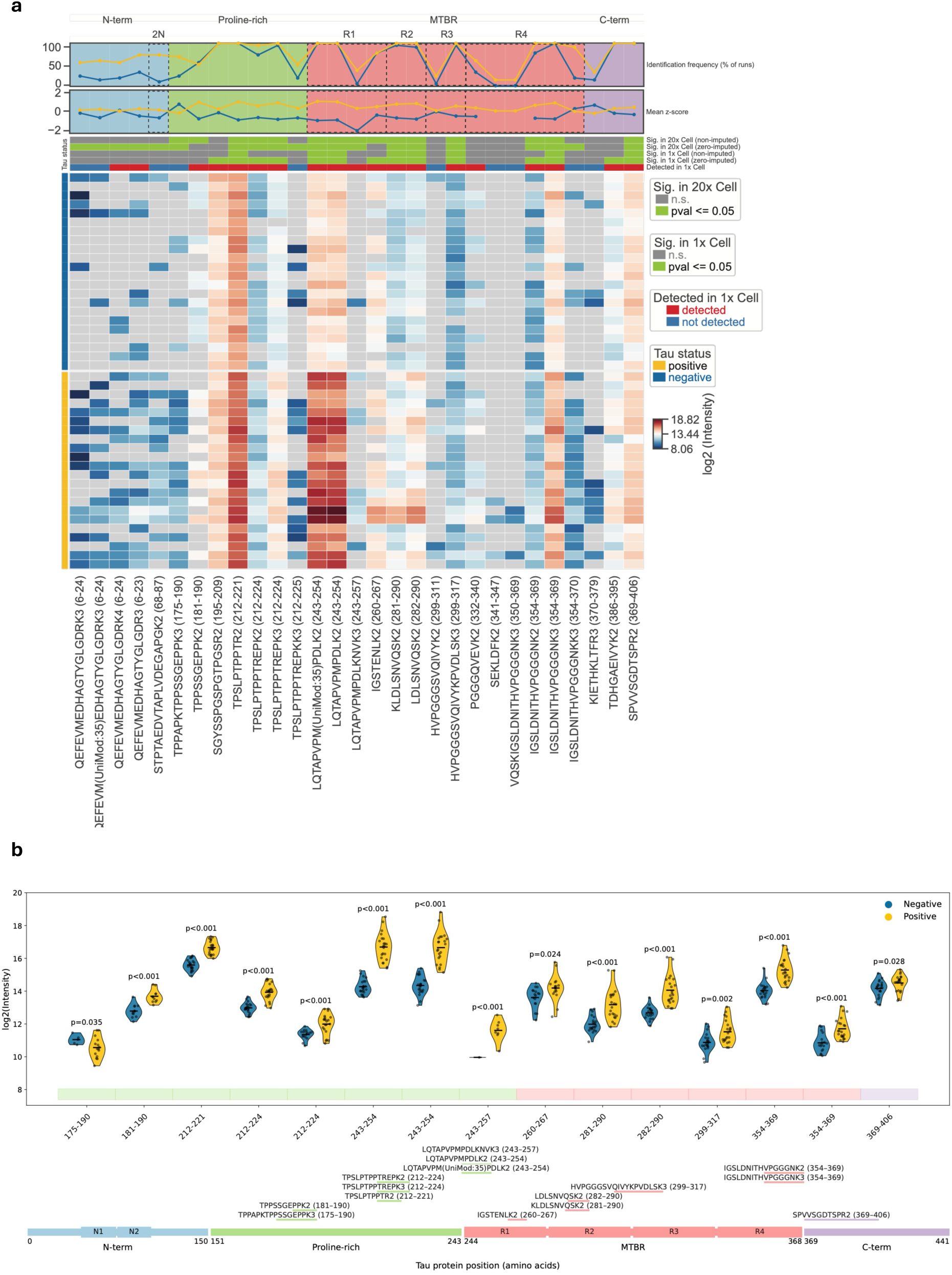
Identified tau peptides. (**a**) Heatmap showing tau protein coverage in positive and negative 20x Cell samples, indicated by the left annotation bars. Rows represent individual measured samples, and columns correspond to individual tau peptides, ordered from N- to C-terminus (left to right). Heatmap colors represent log2 peptide intensities. Grey squares show peptides not detected by LC-MS. Top annotation bars indicate identification of respective peptides in 1x Cell samples and statistical significance (p ≤ 0.05, two-tailed unpaired Welch’s t-test) in 1x and 20x Cell samples, based on non-imputed or zero-imputed data. The line plots above the heatmap show the mean z-score and the frequency of identification of each peptide across samples. (**b**) Violin plots (top) showing intensities of tau peptides significantly different between positive (yellow) and negative (blue) neurons from mini-pool dataset. Points represent individual replicates and lines indicate the mean. The lower panel shows detected peptides mapped onto the tau protein domains. The protein schematic is visually rescaled to expand regions with dense peptide coverage.

**Extended Data Figure 2.**
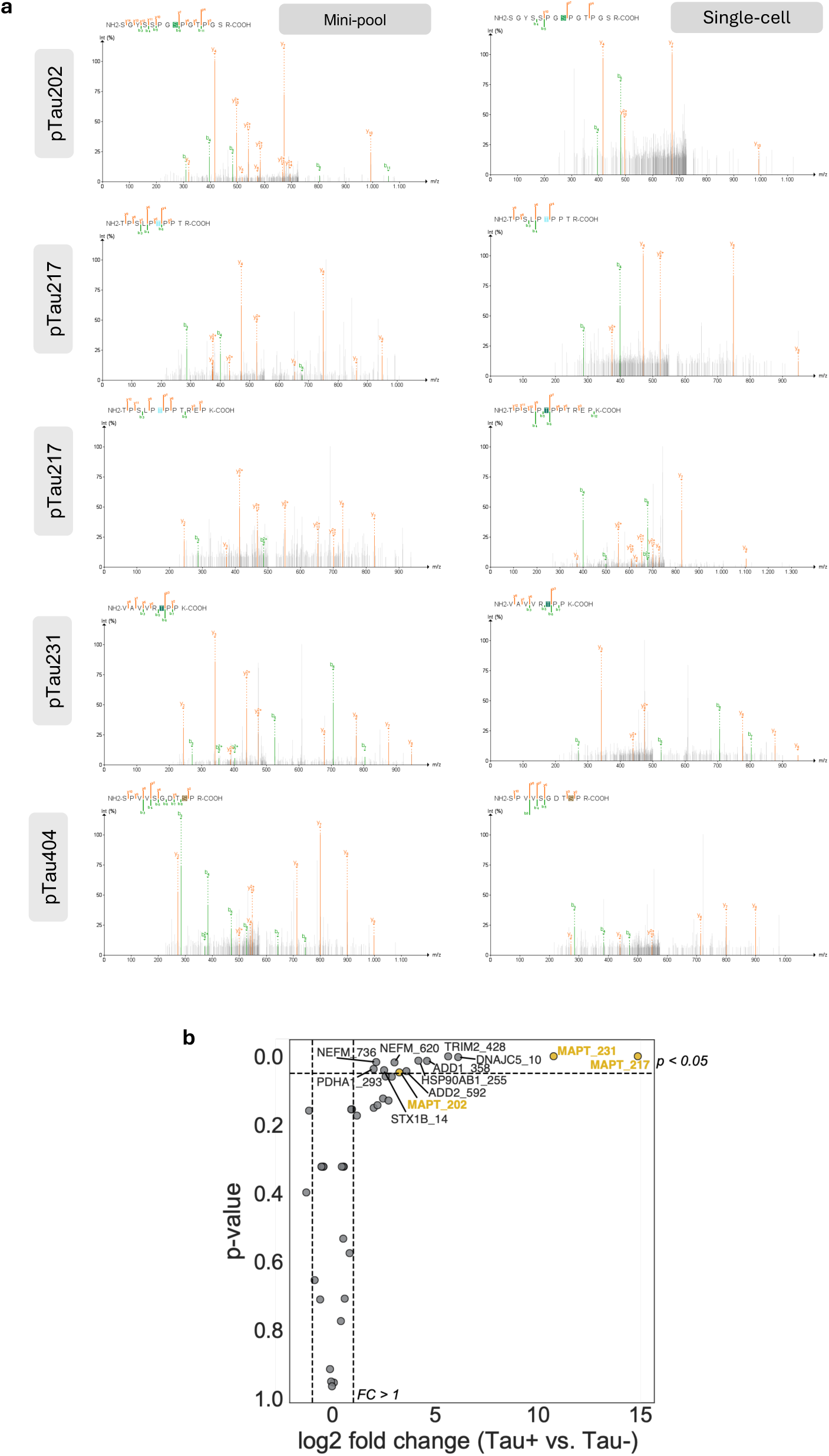
MS spectra for tau phospho-peptides identified. (**a**) Pseudo-MS/MS spectra generated by diaTracer show b- and y-ion series for tau peptides carrying phosphorylation at residues 202, 217, 231 and 404. Left panels correspond to 20x cell samples (mini-pool), right panels to 1x cell samples. Spectra were visualised using the FragPipe-PDV viewer. (**b**) Volcano plot showing phosphorylated protein detected in the mini-pool dataset.

**Extended Data Figure 3.**
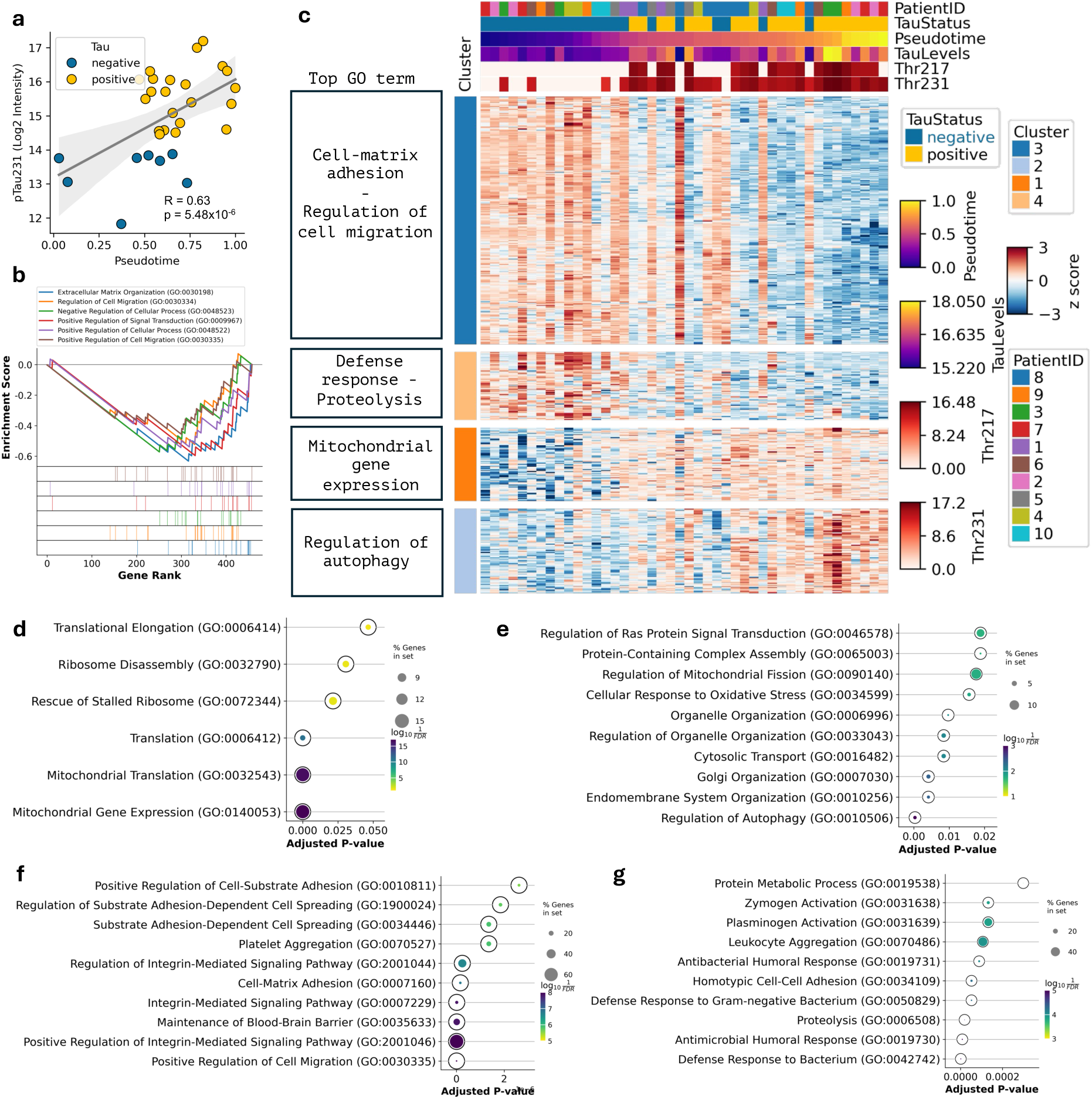
Molecular responses to tangle formation in mini-pool dataset. (**a**) Scatter plot highlighting the positive correlation between Thr231 phosphorylated tau with pseudotime. (**b**) Gene set enrichment analysis for significantly correlated proteins. (**c**) Clustered heatmap of significantly correlated proteins (FDR < 0.05) with pseudotime. Cells (columns) are ordered using pseudotime; pTau status, and Thr217/Thr231 levels and Patient ID are indicated. Top GO terms are shown for each cluster. (**d-g**) GO term analysis for protein in clusters from (c), ranked by adjusted p-value: (d) Cluster 1; (e) Cluster 2; (f) Cluster 3; (g) Cluster 4.

**Extended Data Figure 4.**
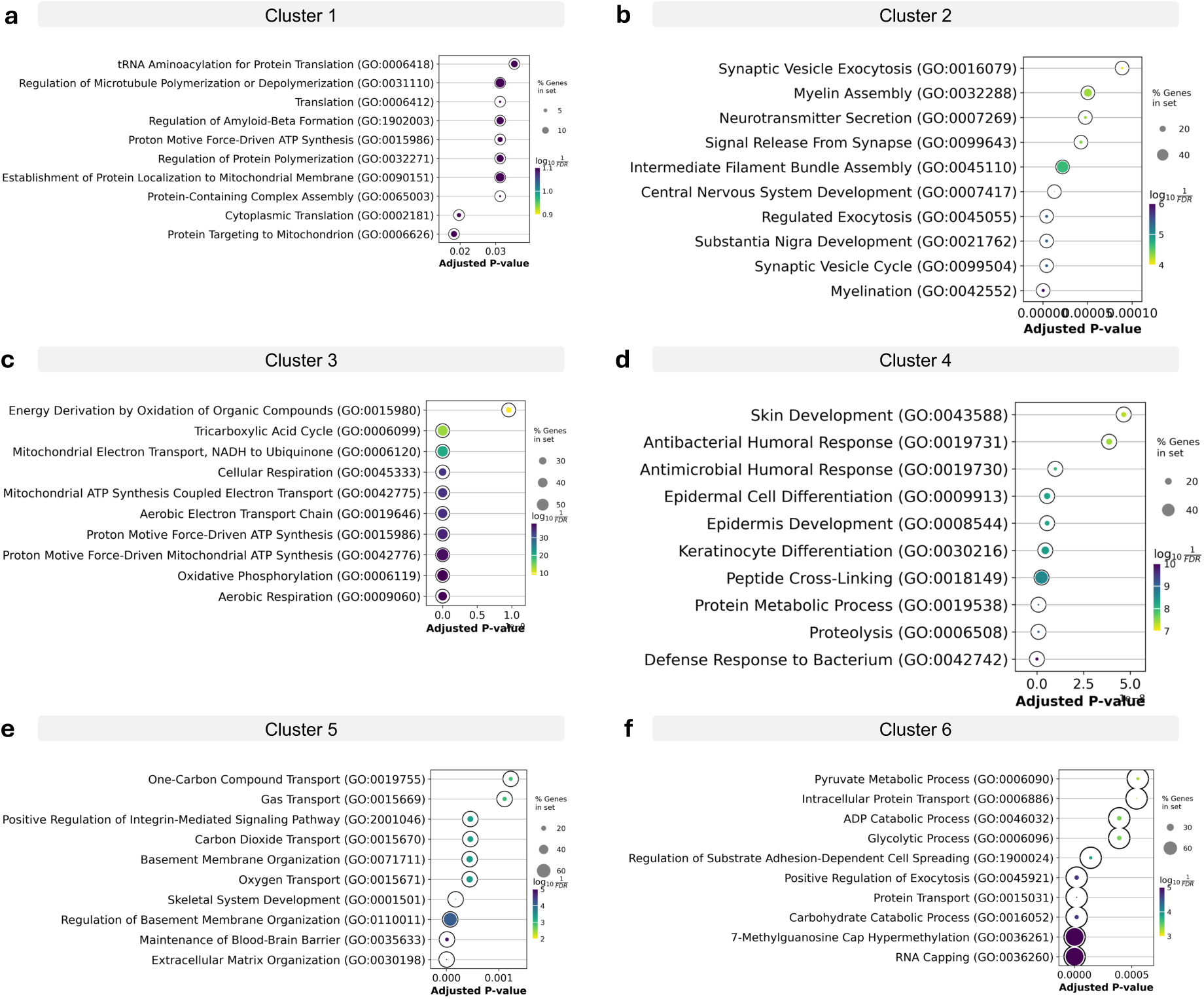
Gene Ontology Analysis of Single-Cell Clusters. GO term analysis for protein in clusters from Fig. 2I, ranked by adjusted p-value: (**a**) Cluster 1; (**b**) Cluster 2; (**c**) Cluster 3; (**d**) Cluster 4; (**e**) Cluster 5; (**f**) Cluster 6.

**Extended Data Figure 5.**
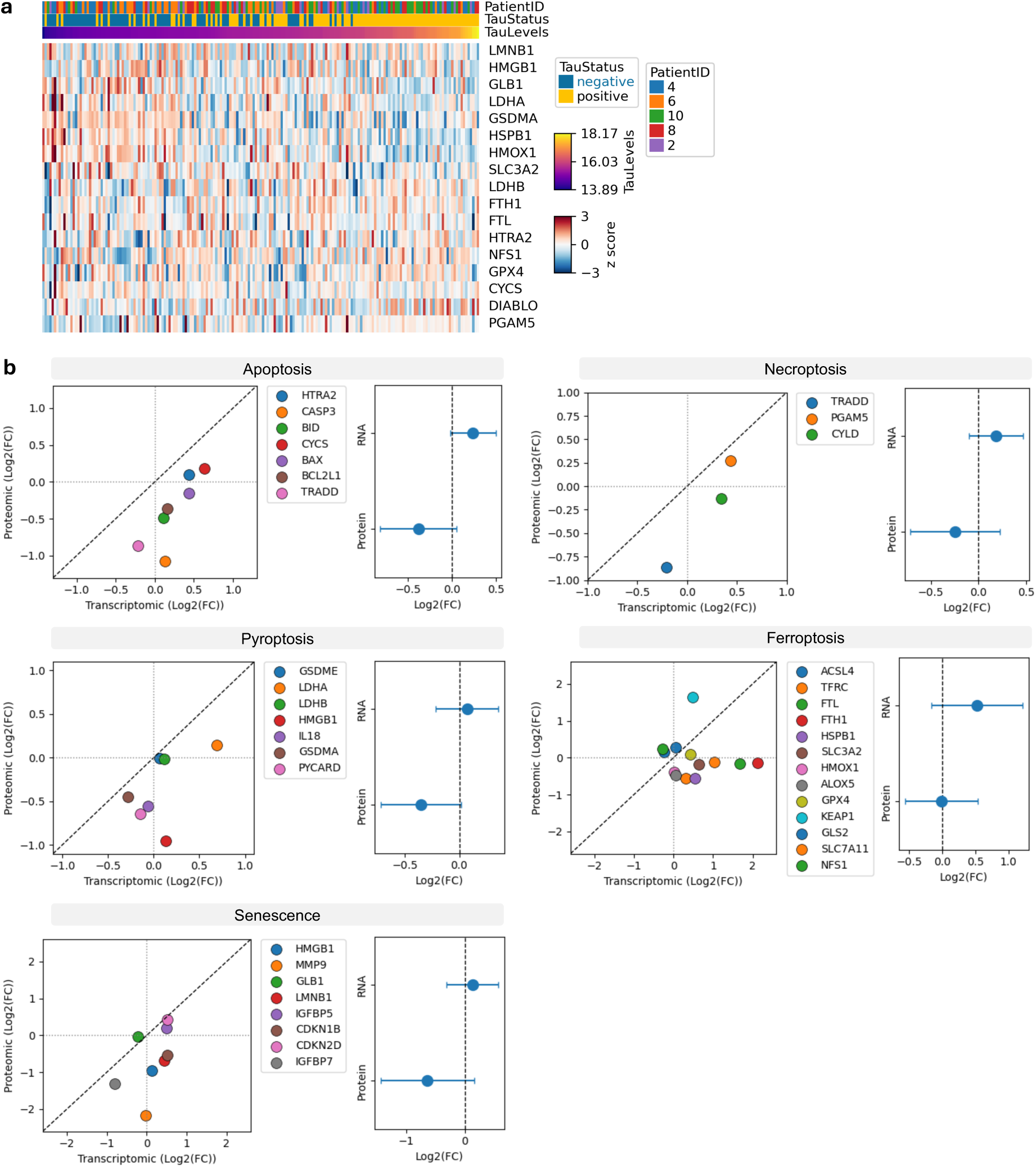
Investigation of cell-death pathways. (**a**) Heatmap of 30 candidate cell-death proteins curated from literature by Edison Analysis AI agent spanning five cell-death pathways (apoptosis, ferroptosis, senescence, necroptosis, and pyroptosis) in the mini-pool dataset. pTau status and Patient ID are indicated. (**b**) Correlation between transcriptomic and proteomic fold changes (log₂FC) for cell-death pathway proteins (left) with summary estimates for both RNA and protein abundance (right). Transcriptome data are from Otero-Garcia *et al*. Neuron 2022.

**Extended Data Figure 6.**
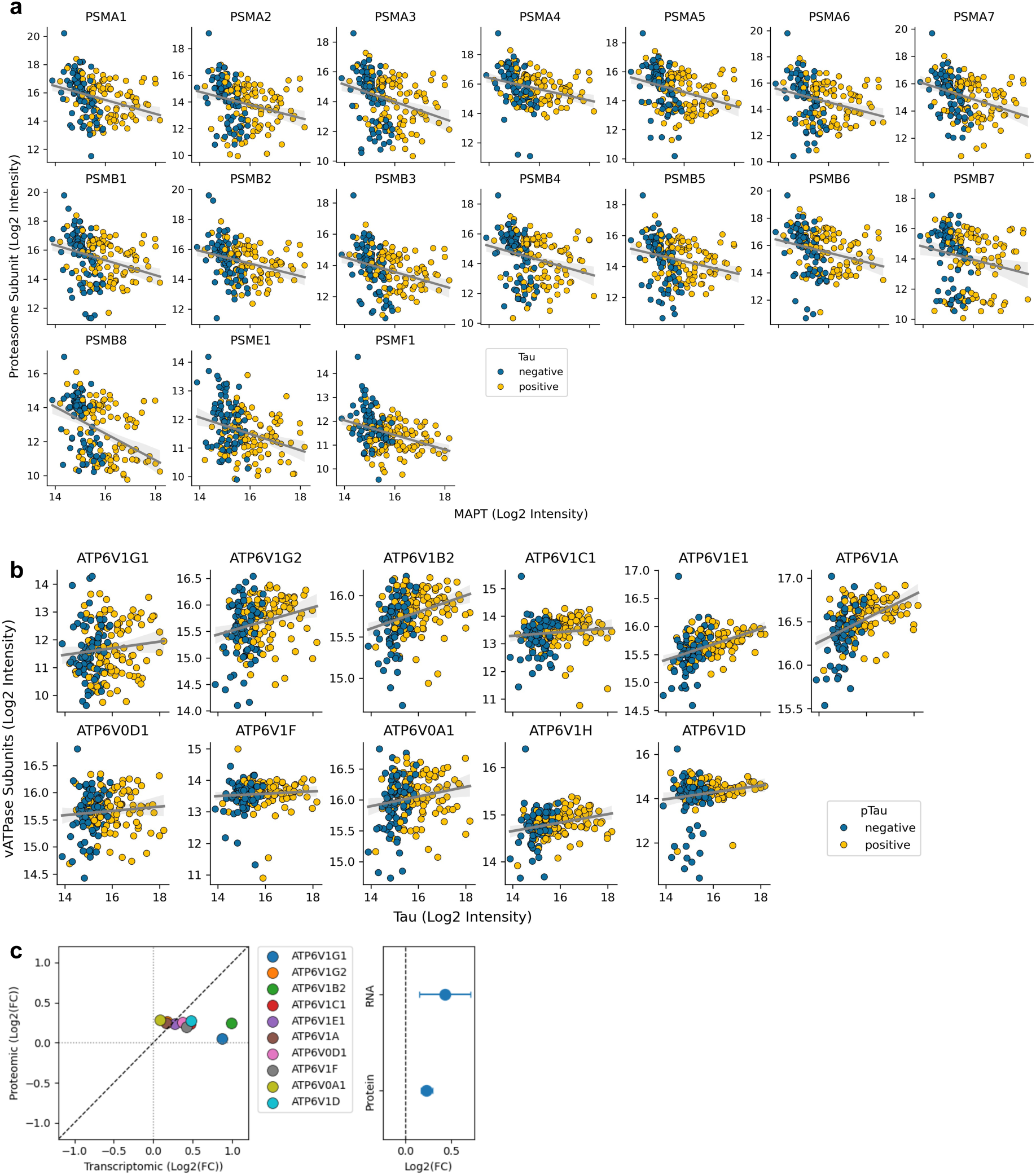
Proteostasis network. (**a**) Proteasome subunit (displaying proteome-wide significant correlation with tau levels) log₂ intensities plotted against total tau (*MAPT*, log₂ intensity) for pTau-negative (blue) and pTau-positive (yellow) cells from the single-cell dataset. (**b**) V-ATPase subunit (ATP6V1 family) log₂ intensities plotted against total tau (log₂ intensity) for pTau-negative (blue) and pTau-positive (yellow) cells from the single-cell dataset. (**c**) Correlation between transcriptomic and proteomic fold changes (log₂FC) for V-ATPase lysosomal subunits (left) with summary estimates for both RNA and protein abundance (right). Transcriptome data are from Otero-Garcia *et al*. Neuron 2022.

